# Effects of N-Acetyl-Leucine and its enantiomers in Niemann-Pick disease type C cells

**DOI:** 10.1101/826222

**Authors:** Danielle te Vruchte, Anthony Galione, Michael Strupp, Michiko Mann

## Abstract

N-Acetyl-DL-Leucine is an acetylated derivative of the essential amino acid leucine and a racemate (1:1) of N-Acetyl-L-Leucine and N-Acetyl-D-Leucine enantiomers. Previous observational clinical studies have demonstrated that N-Acetyl-DL-Leucine is effective in improving ataxia in patients with Niemann-Pick disease type C (NPC), a lysosomal storage disorder characterized at the cellular level by increased relative volume of the endosomal/lysosomal system. In this study, we sought to further characterize the potential therapeutic benefit of N-acetyl-DL-leucine and its enantiomers for the treatment of NPC. We investigated the effectiveness of N-Acetyl-DL-Leucine, N-Acetyl-D-Leucine, and N-Acetyl-L-Leucine in reducing lysosomal volume in non-neuronal NPC1 cells using LysoTracker, a fluorescent dye that accumulates in acidic organelles. We report that both N-Acetyl-DL-Leucine and N-Acetyl-L-Leucine reduced relative lysosomal volume in *NPC1^-/-^* Chinese Hamster Ovary cells in a dose-dependent manner. Subsequently, we confirmed that N-Acetyl-L-Leucine was most effective at reducing relative lysosomal volumes in fibroblasts derived from NPC patients with severe disease (\*\*\**p* <0.001), followed by N-Acetyl-DL-Leucine (\*\**p* <0.01). Treatment with N-Acetyl-D-Leucine did not achieve statistical significance. Taken together, these results suggest that N-Acetyl-L-Leucine is the most effective enantiomer in correcting relative lysosomal volume in non-neuronal NPC cells, and support further research and development of the L-enantiomer.

## Introduction

N-acetyl-DL-leucine is an acetylated derivative of the essential amino acid leucine and a racemic mixture of enantiomers (1:1) N-acetyl-L-leucine and N-acetyl-D-leucine. N-acetyl-DL-leucine has been used in France for over 50 years for the treatment of acute vertigo and has been demonstrated to improve postural compensation in patients after vestibular neurotomy and labyrinthectomy (Leau and Ducrot, 1957; Neuzil, 2002; Viart and Vidal, 2009). A study of N-acetyl-L-leucine and N-acetyl-D-leucine in a rat model of unilateral labyrinthectomy revealed that N-Acetyl-L-Leucine, but not N-Acetyl-D-Leucine, is the pharmacologically active substance that improves central vestibular compensation (Gunther et al., 2015). Furthermore, a study in a unilateral vestibular neurectomy cat model suggested N-Acetyl-L-Leucine is the active part of the racemate that leads to a significant acceleration of the vestibular compensation process, most likely through acting on vestibular nuclei (VN) neurons (Tighilet et al., 2015).

Given that there are phylogenetic and electrophysiological similarities and close interactions between vestibular and deep cerebellar neurons, it was hypothesized that N-Acetyl-DL-Leucine may have clinical utility in the treatment of cerebellar disorders such as Niemann-Pick disease type C (NPC), for example in the reduction of ataxic symptoms, through a potentially similar mechanism observed in models of vertigo. Published studies have shown that this may be because N-acetyl-DL-leucine can correct the membrane potential of hyperpolarized or depolarized vestibular neurons in a lesioned guinea pig model (Vibert and Vidal, 2001), whereby its direct interaction with membrane phospholipids such as phosphatidylinositol 4,5-bisphosphate influenced ion channel activity (Suh and Hille, 2008). Indeed, in observational clinical studies, treatment with N-Acetyl-DL-Leucine improved cerebellar ataxia, including in patients with the lysosomal storage disorder (LSD) NPC) (Strupp et al. 2013; Schneipp et al. 2016; Bremova et al., 2015). However, the pharmacological mechanism of action of N-acetyl-Leucine for these disorders remains to be further clarified.

In this study, we sought to further characterize the potential therapeutic benefit of N-acetyl-DL-leucine and its enantiomers for lysosomal storage disorders such as NPC. In common with other LSDs, NPC is characterized by expansion of the endosomal/lysosomal (LE/Lys) system. Relative volume of the LE/Lys system can be measured in viable cells by flow cytometry using a commercially available fluorescent probe, LysoTracker, a fluorescent dye that is a weak base that can be trapped in the acidic environment of the lysosome (Heard et al., 2010; Lachmann et al., 2004; Wraith, 2002). It has previously been demonstrated that increased LysoTracker fluorescence is a hallmark of NPC1 cells (te Vruchte et al., 2014). Furthermore, this assay has been successfully applied to measure relative acidic compartment volume in B cells from a cohort of NPC patients, as well as *Npc1^-/-^* mutant mice (te Vruchte et al., 2014). This previous study showed that measuring storage in circulating B cells mirrors progressive glycosphingolipid storage in the brain, validating the use of this peripheral cell biomarker for NPC. It is therefore expected that such a biomarker may be broadly used as a universal biomarker for this family of lysosomal storage disorders, as an aid for the initial diagnosis, monitoring disease progression, and determining response to therapies.

Based on the above background, in this study we investigated the impact of N-acetyl-DL-leucine and its enantiomers using LysoTracker in non-neuron cells, including NPC null Chinese Hamster Ovary (CHO) cells and fibroblasts derived from NPC patients. Given that the previous study reported that the N-Acetyl-L-Leucine is the active enantiomer (Gunther et al., 2015; Tighilet et al., 2015), we also compared the effects of N-Acetyl-DL-Leucine, N-Acetyl-L-Leucine and N-Acetyl-D-Leucine on relative lysosomal volume to investigate whether this may also be relevant to the benefits observed in NPC patients.

## Methods

### LysoTracker staining in NPC1 Non-Neuronal Cells

Human fibroblasts from NPC1 patients were purchased from the NINDS Human Genetic Repository at the Coriell Institute (USA). Fibroblasts or wild type (WT) and NPC1-null CHO cell lines cells (Cruz et al., 2000) were grown in T25 or T75 culture flasks (Greiner Bio-One) and treated for 7 days with N-Acetyl-DL-Leucine (Molekula #73891210), N-Acetyl-L-Leucine (Sigma Aldrich #441511) or N-Acetyl-D-Leucine (Sigma Aldrich #A0876) prior to harvesting. Cells were trypsinized, centrifuged (270 x g, 5 min), washed twice with 1x D-PBS (Sigma-Aldrich), centrifuged again and incubated with 100 nM LysoTracker-green DND-26 (Invitrogen) in PBS for 20 min at room temperature. Cells were centrifuged (270 x g, 5 min), resuspended in 0.25 mL of FACS buffer (0.1% BSA, 0.02 M NaN3 in 1 × PBS) and kept on ice for a maximum of 1 h. Prior to flow cytometric analysis (BD Biosciences Accuri C6 Plus), 0.25 mL of FACS buffer with 2 μg/mL propidium iodide (Sigma Aldrich #P4170) was added to the samples. The cytometer was calibrated using Cytometer Setup and Tracking beads (BD #661414), and compensation was performed using cells stained with LysoTracker or propidium Iodide using BD Accuri C6 Plus software (BD). Data were recorded on 10,000 live single cells per sample.

## Results

### Dose-dependent effects of N-Acetyl-DL-Leucine, N-Acetyl-L-Leucine and N-Acetyl-D-Leucine in CHO cells

Wildtype and NPC1-null CHO cells were treated with N-Acetyl-DL-Leucine, N-Acetyl-L-Leucine or N-Acetyl-D-Leucine. CHO cells were incubated with various concentrations (1 μM, 10 μM, 100 μM, 1 mM, 5 mM) of N-Acetyl-DL-Leucine, N-Acetyl-L-Leucine and N-Acetyl-D-Leucine for 7 days, during which the culture medium was refreshed on day 3 or 4. LysoTracker fluorescence was measured on day 7 using flow cytometry and the data was normalized to untreated cells of CHO WT or CHO NPC1. FACS analysis revealed that, whilst differing concentrations of N-Acetyl-DL-Leucine, N-Acetyl-L-Leucine and N-Acetyl-D-Leucine failed to alter the fluorescence signals of LysoTracker in the WT CHO cell line (Figure 1A, B and C, CHO WT in black), fluorescence signals obtained from NPC1-null CHO cells treated with N-Acetyl-DL-Leucine and N-Acetyl-L-Leucine displayed a trend of reduced relative lysosomal volume in a dose-dependent manner (Figure 1A, 5 mM N-Acetyl-DL-Leucine in blue, Figure 1B, 1 mM and 5 mM N-Acetyl-L-Leucine in red), indicating that N-Acetyl-DL-Leucine and N-Acetyl-L-Leucine contributed to reducing LE/Lys volume in this system. On the other hand, N-Acetyl-D-Leucine failed to show a clear trend of dosedependent effects on LysoTracker signals but achieved statistical significance at a higher concentration (Figure 1C, N-Acetyl-D-Leucine at 5 mM in grey).

**Figure 1.**
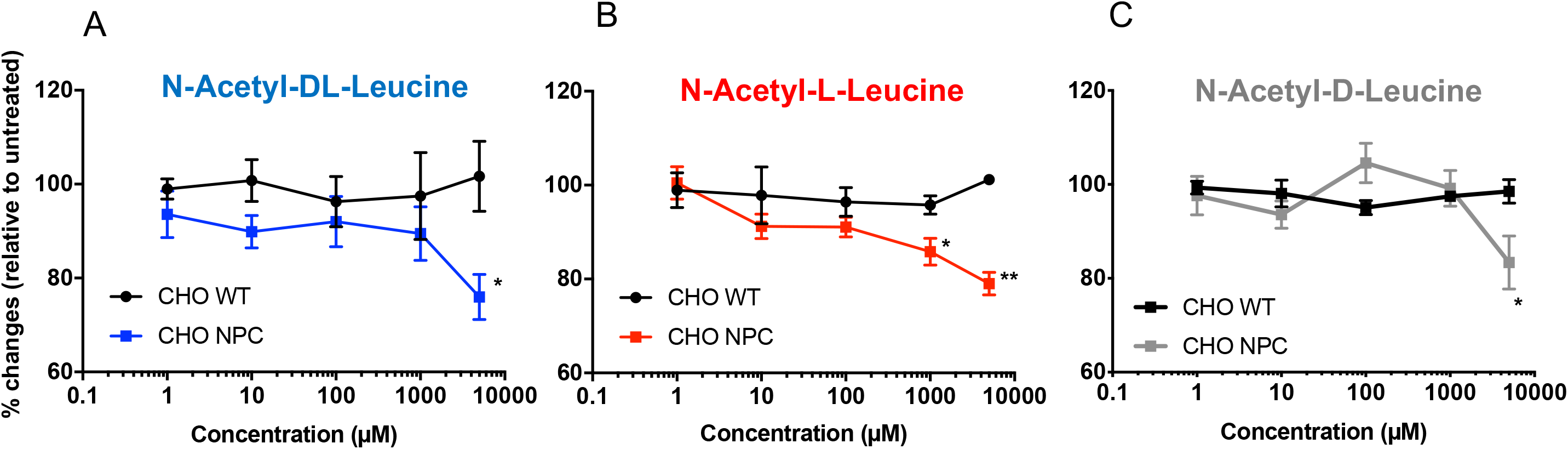
Dose dependent effect of N-Acetyl-DL-Leucine, N-Acetyl-L-Leucine and N-Acetyl-D-Leucine in Wild type CHO and NPC1-null CHO cell lines. Wild type CHO cells (CHO WT) and NPC1-null CHO cells (CHO NPC) were treated with the indicated concentration of **N-Acetyl-DL-Leucine, N-Acetyl-L-Leucine or N-Acetyl-D-Leucine** for 7 days before FACS analysis. Fluorescence signals were normalized to untreated cells (CHO WT or CHO NPC). At each concentration, the values were compared between CHO WT and CHO NPC by conducting an unpaired t test. **(A) N-Acetyl-DL-Leucine treated group (%)** at 5 mM: 101.7 ± 7.4 (CHO WT) and 76.0 ± 4.8 (CHO NPC), * *p* < 0.05. **(B) N-Acetyl-L-Leucine treated group (%)** at 1 mM: 95.8 ± 1.9 (CHO WT) and 85.8 ± 2.9 (CHO NPC), * *p* < 0.05 and at 5 mM: 101.2 ± 1.2 (CHO WT) and 79.0 ± 2.4 (CHO NPC), ** *p* < 0.01. n > 3. **(C) N-Acetyl-D-Leucine treated group (%)** at 5 mM: 98.5 ± 2.5 (CHO WT) and 83.4 ± 5.7 (CHO NPC), * *p* < 0.05. Statistical difference between groups were identified by two-tailed unpaired t test based on n > 3. Mean ± S.E.M.

### N-Acetyl-DL/L/D-Leucine in NPC1 human fibroblasts

Subsequently, we tested if these compounds showed similar responses in human fibroblasts derived from NPC1 patients who displayed severe clinical presentation. Fibroblasts were treated with 1 mM N-Acetyl-DL-Leucine, N-Acetyl-L-Leucine and N-Acetyl-D-Leucine for 7 days, and LysoTracker Green fluorescence signal was determined by FACS. These data were compared with those obtained from an age matched healthy individual as a control. Untreated NPC1 patient fibroblasts exhibited higher values of LysoTracker fluorescence signal (Figure 2A-C) relative to a healthy control, consistent with previous studies in B cells derived from a cohort of NPC1 patients (te Vruchte et al., 2014). Incubating NPC1 patient fibroblasts for 7 days with 1 mM N-Acetyl-DL-Leucine and N-Acetyl-L-Leucine led to a statistically significant decrease in fluorescent signal (Figure 2A and B). Although N-Acetyl-D-Leucine showed a trend of reduced fluorescent signal, it failed to reach statistical significance (Figure 2C).

**Figure 2.**
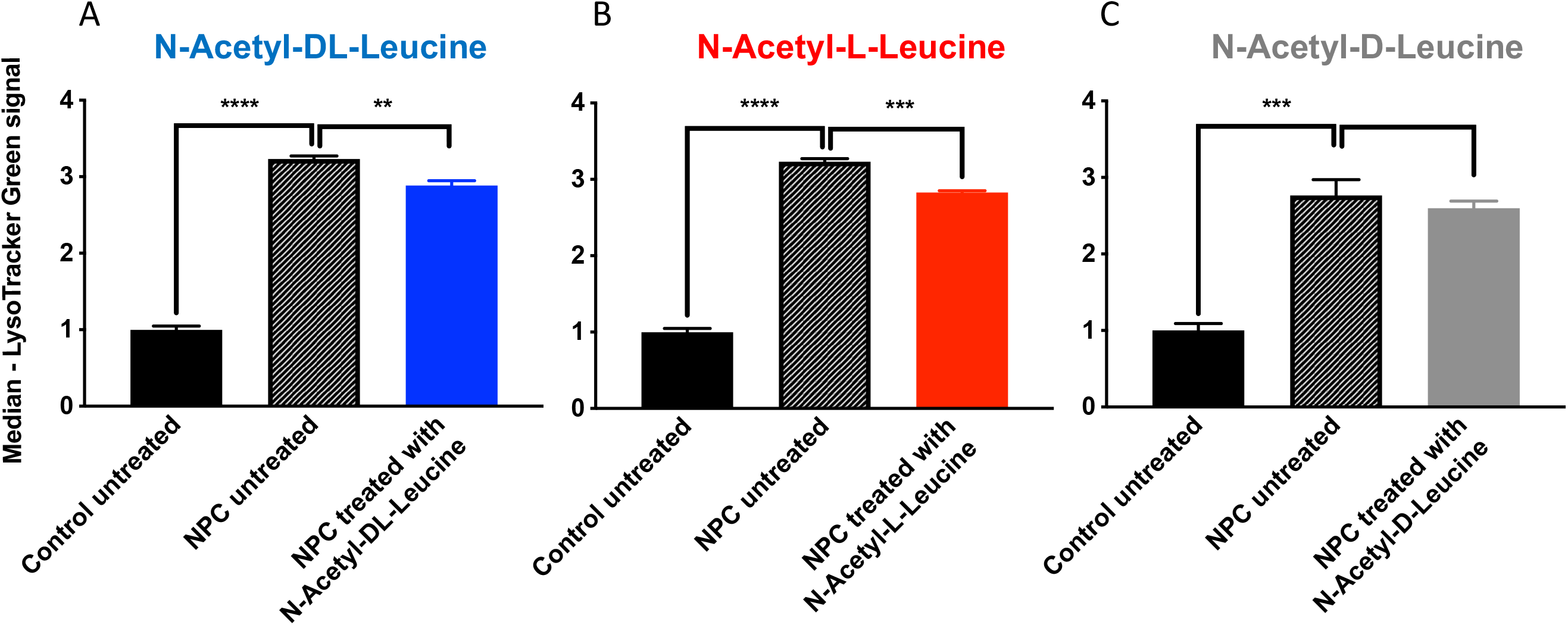
LysoTracker fluorescence in human fibroblasts derive from a healthy age matched individual and NPC1 patients. Cells were treated with 1mM N-Acetyl-DL-Leucine, N-Acetyl-L-Leucine or N-Acetyl-D-Leucine in 0.05% DMSO for 7 days. LysoTracker fluorescence was measured and is expressed as fold change relative to averaged value of untreated control fibroblasts derived from the healthy individual. (**A) N-Acetyl-DL-Leucine treated group;** Control untreated: 1.00 ± 0.05, NPC1 untreated: 3.23 ± 0.04, NPC1 treated with N-Acetyl-DL-Leucine: 2.89 ± 0.06. **(B) N-Acetyl-L-Leucine treated group;** Control untreated: 1.00 ± 0.05, NPC1 untreated: 3.23 ± 0.04 and NPC1 treated with N-Acetyl-L-Leucine: 2.83 ± 0.02. **(C) N-Acetyl-D-Leucine treated group;** Control untreated: 1.00 ± 0.09, NPC1 untreated: 2.77 ± 0.21, NPC1 treated with N-Acetyl-DL-Leucine: 2.60 ± 0.09. **/***/**** - *p* < 0.01/0.001/0.0001, n>3, one-way ANOVA with Tukey’s multiple comparison test was performed. Mean ± S.E.M.

## Discussion and Summary

In the present study, we applied the fluorescent probe LysoTracker to non-neuronal cells, including NPC1-null CHO cells and fibroblasts derived from NPC1 patients, to determine whether N-Acetyl-DL-Leucine, N-Acetyl-L-Leucine and N-Acetyl-D-Leucine would influence LE/Lys compartment. Consistent with the previous report (te Vruchte et al., 2014), we first established that both NPC1-null CHO cells and NPC1 patient fibroblasts exhibited higher lysosomal volumes e relative to controls (Figure 2A-C), representing the enlarged lysosomal storage, a prominent NPC1 cellular phenotype. Second, we found that incubating cells with N-Acetyl-DL-Leucine and N-Acetyl-L-Leucine reduced LysoTracker fluorescence values in a dose-dependent manner in NPC1-null CHO cells (Figure 1 A and B). N-Acetyl-D-Leucine was effective only at the highest concentration (5 mM) (Figure 1C). Finally, when we tested these compounds in the NPC1 patient-derived fibroblasts, we found that N-Acetyl-L-Leucine was most effective in lowering LysoTracker signals followed by N-Acetyl-DL-Leucine (Figure 2).

Our result in non-neuronal cells shows N-acetyl-L-leucine and N-acetyl-DL-leucine have the ability to reduce lysosomal storage in non-neuronal NPC cell types. This is one possible mode of action in LSD which in particular could lead to a systemic disease-modifying effects in these diseases. The results also agree with previous *in vivo* observations suggesting that N-acetyl-L-leucine is the active enantiomer, and which may have superior effects when administered as a single enantiomer (Gunther et al., 2015; Tighilet et al., 2015).

## Notes

**Conflicts of Interest** A.G. is a cofounder, shareholder and consultant to IntraBio. M.Y-M and D.t.V. are shareholders in IntraBio. M.S. acts as a consultant for Abbott, Actelion, AurisMedical, Heel, IntraBio and Sensorion, is a shareholder of IntraBio and Joint Chief Editor of the Journal of Neurology, Editor in Chief of Frontiers of Neuro-otology and Section Editor of F1000. M.S. has received speaker’s honoraria from Abbott, Actelion, Auris Medical, Biogen, Eisai, Gru□nenthal, GSK, Henning Pharma, Interacoustics, Merck, MSD, Otometrics, Pierre-Fabre, TEVA, UCB.

